# Mechanisms of CD4+ T tolerance to a corneal epithelial neoantigen

**DOI:** 10.64898/2026.04.14.718505

**Authors:** Jeremias G. Galletti, Kaitlin K. Scholand, Jianming Shao, Prashant Kumar, Elizaveta A. Demianova, Elle Joy J San Juan, Laura Schaefer, Cintia S. de Paiva

## Abstract

Tissue-specific peripheral tolerance mechanisms are essential to prevent autoimmunity. The cornea is immune privileged, and anterior chamber-associated immune deviation (ACAID) governs its inner surface. However, the mechanisms that apply to corneal epithelial (outer surface) antigens remain unknown. Using an inducible, cornea-restricted neoantigen mouse model, we found that the cornea relies on inducible regulatory T cells (Tregs) rather than ignorance or ACAID for its epithelial antigens. Although the cornea is both avascular and alymphatic, its epithelial antigens are still efficiently presented by ocular surface-derived antigen-presenting cells to T cells in draining lymph nodes under homeostatic conditions, leading to conventional antigen-specific Treg expansion without ocular pathology. This tolerance was not absolute: systemic immunization redirected antigen-specific responses toward pathogenic effector T cells that disrupted epithelial barrier function. These findings identify Treg induction as a dominant mechanism of corneal epithelial immune homeostasis and demonstrate that inflammatory priming can render a tolerated corneal antigen into an autoimmune target, providing mechanistic insight into dry eye pathogenesis.

**Summary:** This study shows that immune tolerance to corneal epithelial neoantigens relies not on immune privilege but on peripherally induced regulatory T cells in the draining lymph nodes that can be subverted by innate activation, shedding light on ocular surface disease pathophysiology.

## Introduction

The cornea is an avascular, transparent tissue that focuses light rays on the retina, but only when its surface is kept moist and smooth by the tear film. Immune cells such as dendritic cells, macrophages, and γδ T cells are present in small numbers in the cornea (Alam et al., 2024; Downie et al., 2023; Loi et al., 2022; Wu et al., 2024) and participate in immune surveillance. However, the cornea must stay uninflamed to maintain clarity (Downie et al., 2021:Chen, 2022 #9825). The ocular surface has evolved into a mucosal environment enclosed within the eyelids, providing the cornea with protection and a healthy tear film, and thus, enabling proper sight (Galletti et al., 2017). Conversely, a dysfunctional tear film (in quantity or quality) is the defining feature of the prevalent ocular surface disorder known as dry eye disease (Craig et al., 2017). This condition greatly impacts patients’ quality of life, and its incidence is rising due to environmental factors (Stapleton et al., 2017).

Autoreactive CD4^+^T cells drive corneal and conjunctival inflammation in dry eye (Chen et al., 2014; Niederkorn et al., 2006; Vereertbrugghen et al., 2024). Since the cornea lacks blood vessels, local T cell activity depends on limbal and conjunctival T cells extravasating to these tissues. Although some CD4^+^T cells enter and actively patrol the corneal epithelium (Downie et al., 2023; Loi et al., 2022; Wu et al., 2024), most stay in the neighboring limbus and conjunctiva, from where they exert their effects on the corneal effects (Vereertbrugghen et al., 2024). In dry eye, ocular surface desiccation damages the corneal epithelium and activates innate immunity, leading to antigen presentation and differentiation of naïve T cells into effector subsets such as Th1 and Th17 cells (Bose et al., 2017; Bron et al., 2017; Foulsham et al., 2017; Stern et al., 2002). These cells infiltrate ocular surface tissues, release pro-inflammatory cytokines, and perpetuate epithelial damage, driving the chronic inflammatory cycle underlying dry eye (Bron et al., 2017). The resulting corneal epitheliopathy leads to dry eye symptoms such as blurry sight, ocular discomfort, and eye pain (Bron et al., 2017; Tsubota et al., 2020).

To prevent unwanted inflammation, the immune system must discriminate between self and non-self to eliminate pathogens while preserving host tissues (Meng et al., 2023). Autoreactive T cells are normally controlled through central and peripheral tolerance. Central tolerance occurs in the thymus, where strongly self-reactive T cells are deleted or become regulatory T cells (Tregs)(Salaman and Gould, 2020). However, thymic expression of self-antigens is incomplete (Danan-Gotthold et al., 2016) and is supplemented by additional peripheral tolerance mechanisms which are tissue-specific (Legoux et al., 2015; Malhotra et al., 2016). Regarding the eye, several peripheral tolerance mechanisms have been described (Galletti and de Paiva, 2021; Galletti et al., 2017; Vendomèle et al., 2017). First and foremost is anterior chamber-associated immune deviation (ACAID), a unique systemic antigen-specific immunosuppressive pathway that underpins the cornea’s immune privilege and enables its transplantation without tissue typing (Chen et al., 2022; Hori et al., 2019; Taylor, 2016). This mechanism, however, involves antigen exposure in the anterior chamber of the eye and thus applies only to the corneal endothelium that lines the inner corneal surface. By contrast, the corneal epithelium is exposed to the environment and exhibits other distinctive immunoregulatory mechanisms, including constitutive expression of PD-L1, FasL, TSP-1, and VEGFR1-3, all of which contribute to allospecific tolerance (Chen et al., 2022; Hori et al., 2019). Ocular exposure to an exogenous antigen leads to systemic tolerance via a process termed mucosal tolerance, which involves antigen uptake by conjunctival antigen-presenting cells and the generation of Tregs (Galletti and de Paiva, 2021; Guzman et al., 2014; Ko et al., 2018). Despite numerous attempts, the targets of autoreactive T cells in dry eye remain unidentified (Bacman et al., 2001; Baer and Hammitt, 2021; Cottin, 2022; El Annan et al., 2013; Kuwana et al., 2001; Moustardas et al., 2021; Reina et al., 2011; Szczerba et al., 2016; Wu et al., 2015). Corneal epithelial autoantigens are likely involved, as these cells sustain immune-mediated damage in this disease (Chen et al., 2014; Niederkorn et al., 2006; Vereertbrugghen et al., 2024). Nonetheless, the tolerance mechanisms concerning antigens expressed directly within the corneal epithelium remain poorly understood, and have only been explored indirectly in the setting of allogeneic corneal transplantation (Chen et al., 2022; Hori et al., 2019).

In the periphery, a naïve autoreactive T cell may undergo deletion, anergy, or Treg differentiation upon encountering its cognate autoantigen, or remain immunologically ignorant. The outcome depends on the tissue involved, antigen availability, and the inflammatory context (Legoux et al., 2015; Malhotra et al., 2016; Salaman and Gould, 2020). Environmental stressors such as pollution, microbial products, and low humidity can activate innate immune signaling in the ocular surface epithelium, potentially breaking tolerance and enabling pathogenic T-cell responses (Guzman et al., 2016a; Guzman et al., 2016b; Guzmán et al., 2020; Guzman et al., 2014). Therefore, understanding tolerance to corneal epithelial antigens is critical for explaining how ocular surface immune homeostasis is maintained and how it fails in disease (Galletti and de Paiva, 2021). However, studying these mechanisms in a non-allogeneic corneal transplantation context is more challenging because antigen-specific CD4^+^T cells represent only a small fraction of the total T cell population (<1–2%) (Bacher and Scheffold, 2013), and tracking them requires prior knowledge of the antigen. To address this, we first developed an inducible mouse model of corneal epithelium-restricted ovalbumin expression, which allowed us to delineate the tolerance pathways involved and the fate of self-reactive CD4⁺T cells specific for corneal epithelial antigens.

## Results

### Development and validation of a cornea-restricted neoantigen model

Tissue-restricted expression of a genetically introduced antigen in otherwise normal mice allows the detailed investigation of tolerance and autoimmunity mechanisms that are organ-specific (Cebula et al., 2013; Legoux et al., 2015; Riehn et al., 2017; Salaman and Gould, 2020). Since a cornea-restricted system was lacking, we first developed a mouse model that expressed an ovalbumin fragment (OVA) as a self-antigen exclusively in the corneal epithelium. To this aim, we combined the ROSA26OVA^f/f^ mice (Cebula et al., 2013; Legoux et al., 2015; Riehn et al., 2017), which harbor a reversed (transcriptionally inactive) OVA sequence flanked by opposing LoxP sites, with keratin 12-reverse tetracycline-trans-activator mice (Chikama et al., 2005) and tetOCre mice (**Fig. 1A**). This resulted in a ternary mouse model that expressed OVA only in corneal epithelial cells upon Cre recombinase induction through tetracycline-containing chow (**Fig. 1B**). Thus, from an immunological viewpoint, OVA represented a corneal epithelial neoantigen (not expressed until induced) in this mouse.

**Figure 1.**
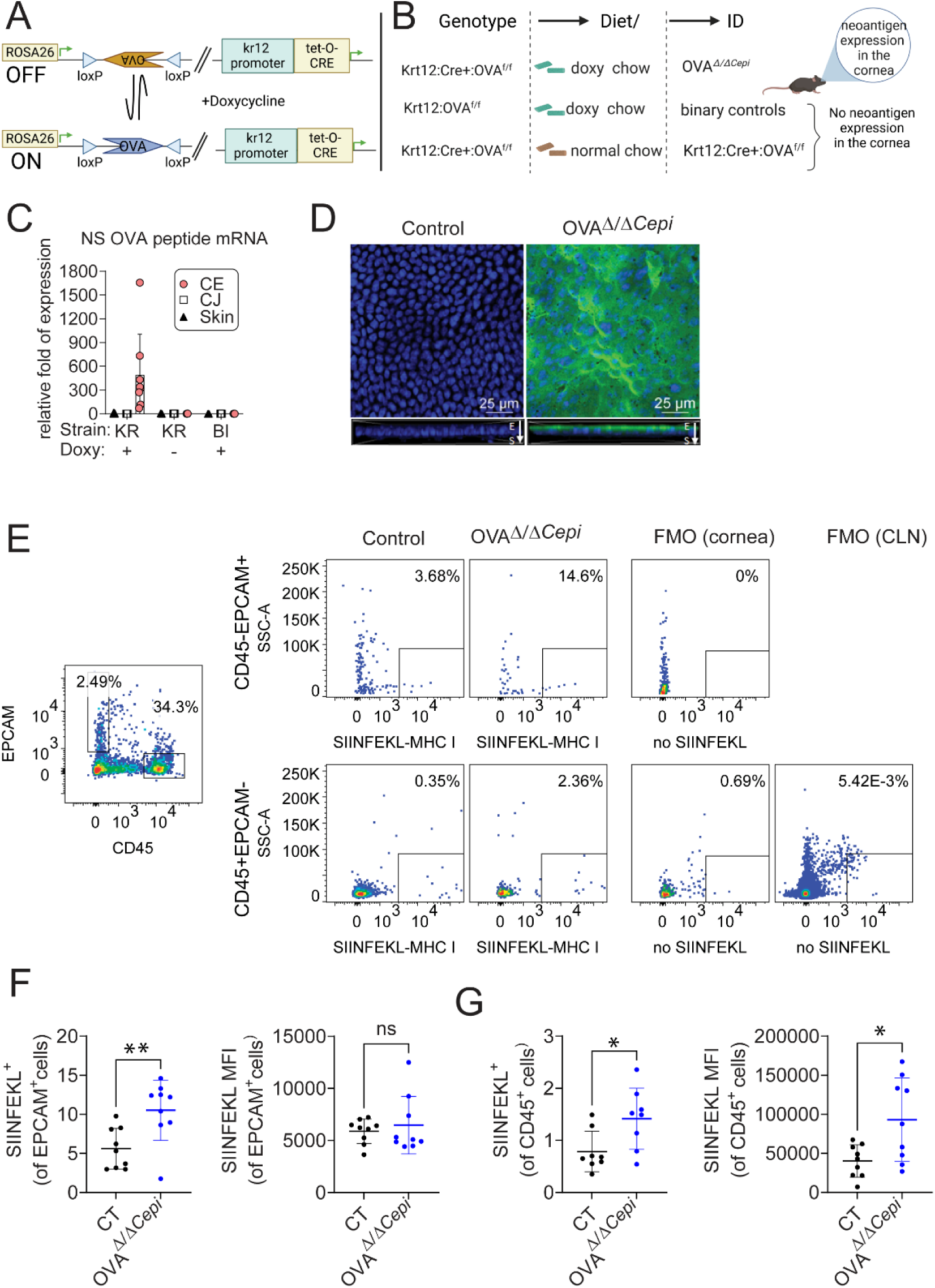
Transgenic mouse model validation. **A.** Schematic of ON and OFF OVA with and without doxycycline diet. Modified from Cebula et al. **B.** Possible genetic and diet combination. Neoantigen expression is expected only in the **OVA*^Δ/ΔCepi^*** mice. **C. OVA mRNA is only expressed in the naïve OVA*^Δ/ΔCepi^* corneal epithelium**, qPCR using specific primers for OVA fusion protein, n = 4-9/group. BI = binary mouse (Kr12:OVA^f/f^); CO = cornea; CJ = conjunctiva; Doxy = doxycycline chow; KR = Krt12:Cre+:OVA^f/f^. OVA*^Δ/ΔCepi^* are KR mice receiving doxycycline chow. **D.** Merged images (en face) of laser confocal microscopy of whole-mount corneas stained with anti-OVA antibody (green) with DAPI nuclear staining (blue). Insets show reconstructed cross-sections using the Z-stack function. E = epithelium; S = stroma **E.** Representative flow plots of single-cell cornea suspensions stained with EPCAM, CD45, and SIINFEKL-MHC I conjugated antibodies. **F-G.** Cumulative data of frequency (**F**) and median fluorescence intensity (**G**). Each dot represents an individual mouse. Mann-Whitney *U* test. CT = control. All mice received doxycycline for at least 2 weeks. *P<0.05; **P<0.01, ns = non-significant

First, we confirmed the cornea-restricted OVA expression of the model using qPCR and immunostaining (Cebula et al., 2013; Ehst et al., 2003; Riehn et al., 2017). After dietary induction for 2 weeks, we compared OVA mRNA expression in the corneal epithelium, conjunctiva, and skin between OVA*^Δ/ΔCepi^* (our model) and cage mate control mice. We included Krt12:Cre^+^:OVA^f/f^ without doxycycline chow to rule out off-target OVA expression due to ectopic Cre recombination(Wu et al., 2020). Only OVA*^Δ/ΔCepi^* corneal epithelium had OVA transcripts **(Fig. 1C),** confirming the specificity of our model. Validating the qPCR results, only the OVA*^Δ/ΔCepi^* mice showed OVA protein immunoreactivity in the corneal epithelium (**Fig. 1D**). Next, we verified that the newly expressed OVA was handled as other endogenous proteins by corneal epithelial cells. Since OVA was expressed as a cytosolic protein in this model, we expected corneal epithelial cells to process and present OVA-derived antigenic peptides on their MHC class I molecules. To verify this, we used the extensively validated 25-D1.16 monoclonal antibody, which only binds to OVA-derived SIINFELK peptide-MHC I complexes on the cell surface. As shown in **Fig. 1E** and **1F**, corneal epithelial (EpCAM^+^) cells from induced OVA*^Δ/ΔCepi^* mice expressed significantly more OVA peptide-presenting MHC I molecules on their surface. We also found a significant increase in the proportion of bone marrow-derived (CD45^+^) corneal cells displaying this peptide on their MHC I molecules (**Fig. 1G**), which is consistent with cross-presentation by antigen-presenting cells after endocytosis of corneal epithelial cell-derived material. Collectively, these results show that in this mouse model, cornea-restricted expression of OVA can be induced and leads to the physiological presentation of its antigenic peptides by corneal epithelial and antigen-presenting cells.

### Eye-draining cells can process and present the cornea-restricted antigen to CD4^+^T cells

APCs orchestrate adaptive immune responses, and their interaction with CD4^+^T cells via MHC II-restricted presentation is a decisive step in antigen-specific peripheral tolerance. To validate our system further, we investigated whether APCs from the OVA^Δ/ΔCepi^ mice were taking up the corneal epithelium-restricted antigen, and thus, whether they were potentially capable of activating OVA-specific CD4^+^T cells. To this aim, we performed a functional assay by co-culturing mitomycin-treated cervical lymph node cell suspensions (as a source of eye-derived APCs) with labeled OVA-specific (OT-II) CD4^+^T cells over 4 days (**Fig. 2A**) and measured their proliferative response using flow cytometry (Ehst et al., 2003). APCs from OVA*^Δ/ΔCepi^* mice effectively presented cornea-derived OVA peptides on their MHC-II molecules, as evidenced by a significant increase in the proliferating CD4^+^T cells over the basal, non-OVA-specific response in control cultures (**Fig. 2B-C**). When soluble OVA was added to cultures from the start as a positive control, CD4^+^T cell proliferation increased with both APC suspensions but still remained higher in OVA*^Δ/ΔCepi^* cultures. This indicates that despite the relatively high expression levels of corneal OVA, its presentation was not saturating the eye-derived APCs. This was consistent with physiological antigen presentation, where only a minor fraction of MHC II molecules are occupied by a specific peptide (Stern and Santambrogio, 2016). Altogether, our results confirm that the OVA^Δ/ΔCepi^ mouse is valid for studying CD4^+^T cell-mediated tolerance and autoimmunity against a corneal antigen.

**Figure 2.**
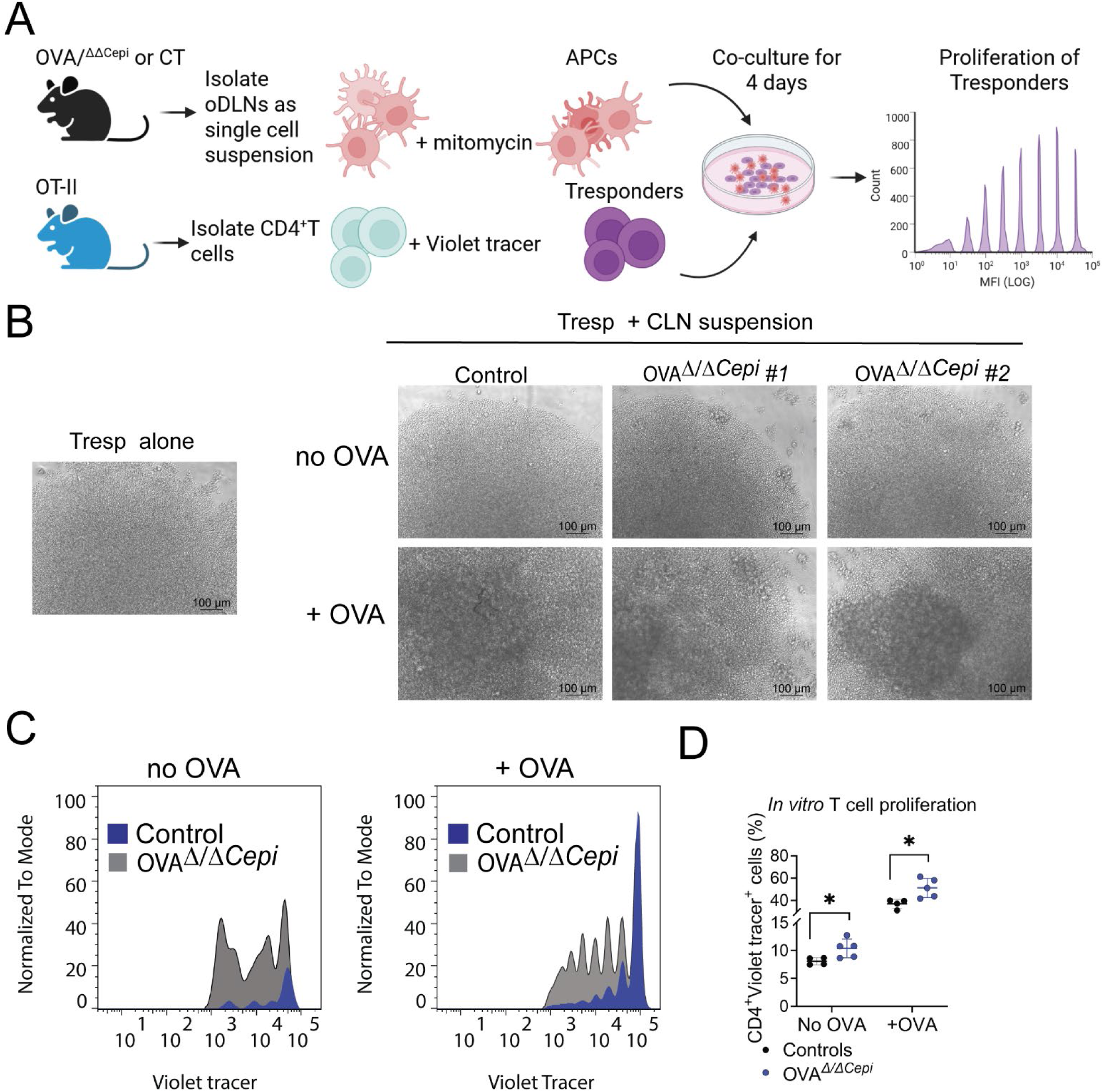
APCs can respond to neoantigen in the cornea, inducing T cell proliferation. Ocular draining lymph nodes from control (CT) and OVA*^Δ/ΔCepi^* mice were excised, filtered, and prepared as a single-cell suspension and incubated with mitomycin C. All mice received doxycycline chow for at least 2 weeks prior to experiments. CD4^+^T cells were isolated from OT-II splenocytes, labelled with Violet Tracer and used as T responders (Tresp). Cervical lymph node (CLN) suspensions and Tresp were mixed *in vitro* with and without exogenous ovalbumin (OVA) and cultured for 4 days. Cell proliferation was performed using flow cytometry and the percentage of proliferating cells was calculated. **A.** Experimental schematic. **B.** Representative bright field images of day 4 cultures of either Tresp alone or Tresp + CLN from the different groups used as APCs. **C.** Representative histograms showing control (gray) and OVA*^Δ/ΔCepi^* (black) without and with OVA. **D.** Frequency of proliferating OTII CD4^+^T Violet Tracer^+^ cells after *in vitro* proliferation assay without and with exogenous OVA. CT = control. Mann-Whitney *U* test. *P<0.05, n = 4-5 animals/group.

### Neoantigen expression is tolerated and does not cause spontaneous dry eye in this model

Immune tolerance is not the mere absence of an immune response but the active result of several regulatory mechanisms. Since no danger signals were present in the ocular surface, we hypothesized that *de novo* corneal OVA expression would be <u>tolerated</u>. To test this, we measured corneal barrier function (as in (Schaefer et al., 2022; Yu et al., 2022)) and cornea mechanosensitivity (as in (Galletti et al., 2023)) after 2 weeks of dietary induction because the cornea epithelium renews itself every 7-10 days. Loss of mechanical sensitivity and epithelial barrier dysfunction are signs of corneal disease in aqueous-tear-deficient dry eye (Adatia et al., 2004; Bourcier et al., 2005; de Paiva et al., 2006; Galletti et al., 2023; Schaefer et al., 2022; Stepp et al., 2018). OVA*^Δ/ΔCepi^* mice had corneal mechanosensitivity (**Fig. 3A**) and corneal barrier levels similar to control mice (**Fig. 3B, 3C**) that was within the normal values obtained in naïve C57BL/6J mice in our publications (Abu-Romman et al., 2024; Yu, 2022; Yu et al., 2022). Because aging can lead to spontaneous dry eye (McClellan et al., 2014), we subjected two-month-old mice to a doxycycline diet for 4 months to investigate if continuous OVA expression in the cornea would lead to spontaneous dry eye. We investigated corneal barrier disruption at 6 months of age, when it physiologically occurs (Yu et al., 2021b). Despite prolonged corneal antigen expression, OVA*^Δ/ΔCepi^* mice had comparable age-related corneal barrier disruption to age-matched controls (**Fig. 3B**). This confirms that corneal OVA expression in this model serves as a surrogate for physiological epithelial antigens, as the age-related mechanisms leading to dry eye are not exacerbated by the transgene (OVA) expression.

**Figure 3.**
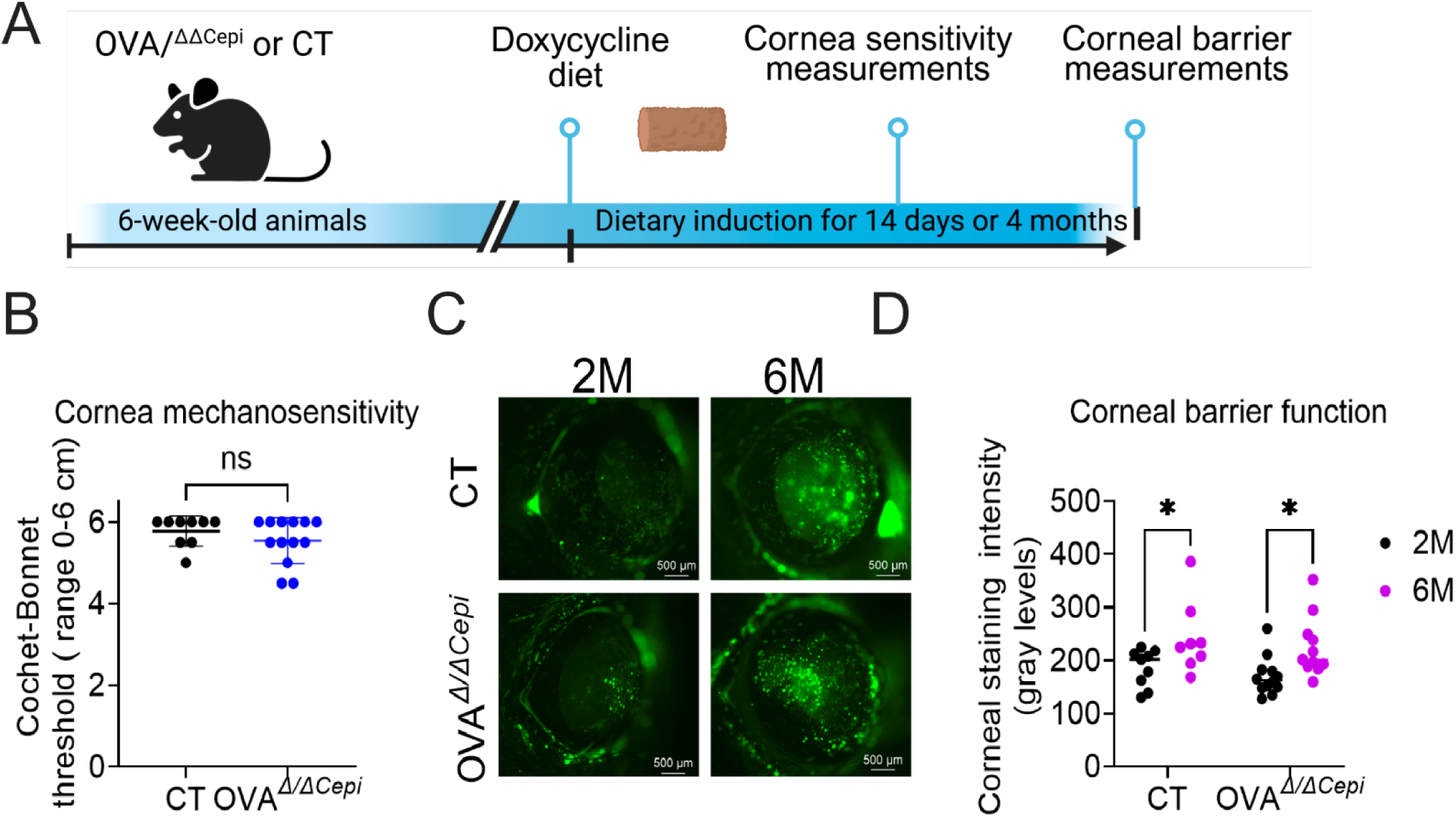
Neoantigen expression does not cause dry eye in naïve conditions. 6-week-old animals control (CT) and OVA*^Δ/ΔCepi^* received doxycycline chow to induce neoantigen expression in the cornea for either 2 weeks or 4 months (M). **A.** Mechanosensitivity measured with Cochet-Bonnet esthesiometer 2 weeks after dietary induction (range 0-6, n = 8-10/group). High values are expected in naive mice and indicate normal range. **B-C**. Corneal barrier function is measured as an uptake of a fluorescent dye at 2 months (M) or 6M (after 2 weeks or 4M post doxycycline diet, respectively). High levels of dye uptake indicate corneal barrier disruption. Representative images are shown in **B** and cumulative data in **C**. There are no interstrain differences at 2M and 6M. CT = control. Mann-Whitney *U* test. *P<0.05, ns = non-significant, n = 4-5 animals/group.

Our previous results (Fig. 2) ruled out antigen sequestration (immunological ignorance) as a tolerance mechanism. Therefore, we hypothesized that immune homeostasis in our system was based on antigen presentation in the absence of ocular surface-derived danger signals. To test this, we examined the number and phenotype of APCs in the eye-draining lymph nodes after corneal OVA induction, which were not expected to be affected by *de novo* autoantigen expression. We used CD86 as an activation marker and IL-12 as a polarizing Th1 cytokine involved in dry eye (Chen et al., 2017). After doxycycline induction, both OVA*^Δ/ΔCepi^* and control mice had similar numbers of conventional dendritic cells (CD45^+^CD11b^+^CD11c^+^MHC II^+^), and within this population, comparable proportions of CD86^+^ and IL-12-producing cells (Supplemental Fig. 1). Overall, these results support our hypothesis that active mechanisms of CD4^+^T cell tolerance prevent the development of a clinical dry eye phenotype, and that this mouse model presents an ideal opportunity to study them in both health and disease.

### OVA-specific T cells encounter and tolerate the corneal antigen in this model mostly by becoming Tregs

Upon neoantigen expression in a tissue, the cognate self-reactive T cells must either become Tregs, anergic or remain immunologically ignorant to avoid autoimmunity. To answer this, we examined the phenotype of corneal antigen-specific T cells in our model. Because antigen-specific T cells are scarce in the physiological T cell repertoire, we increased the experimental power of the system by adoptively transferring 2 × 10^6^ splenic CD4^+^T cells from OT-II mice into OVA*^Δ/ΔCepi^* or control mice which had been on doxycycline chow for two weeks **(Fig. 4A)**. The adoptive transfer of OVA-reactive T cells increased the number of corneal antigen-specific T cells to ease tracking and analysis, but it could also disrupt tolerance and lead to tissue damage by enlarging the autoreactive population. To rule this out, we measured corneal mechanical sensitivity and the ocular response to capsaicin because we have previously shown that these readouts are the most sensitive indicators of T cell-driven corneal tissue damage (Pizzano et al., 2024; Pizzano et al., 2026; Vereertbrugghen et al., 2023). Similar to the lack of dry eye phenotype observed in naïve mice, OT-II adoptively transferred OVA*^Δ/ΔCepi^* recipient mice (with an enlarged corneal antigen-specific T cell pool) did not show a decrease in corneal mechanical sensitivity (**Fig. 4B**) or an increased response to capsaicin (**Fig. 4C-D**) compared to control mice. Corneal barrier function was not measured because the fluorescent dye could interfere with the flow cytometry experiments described below.

**Figure 4.**
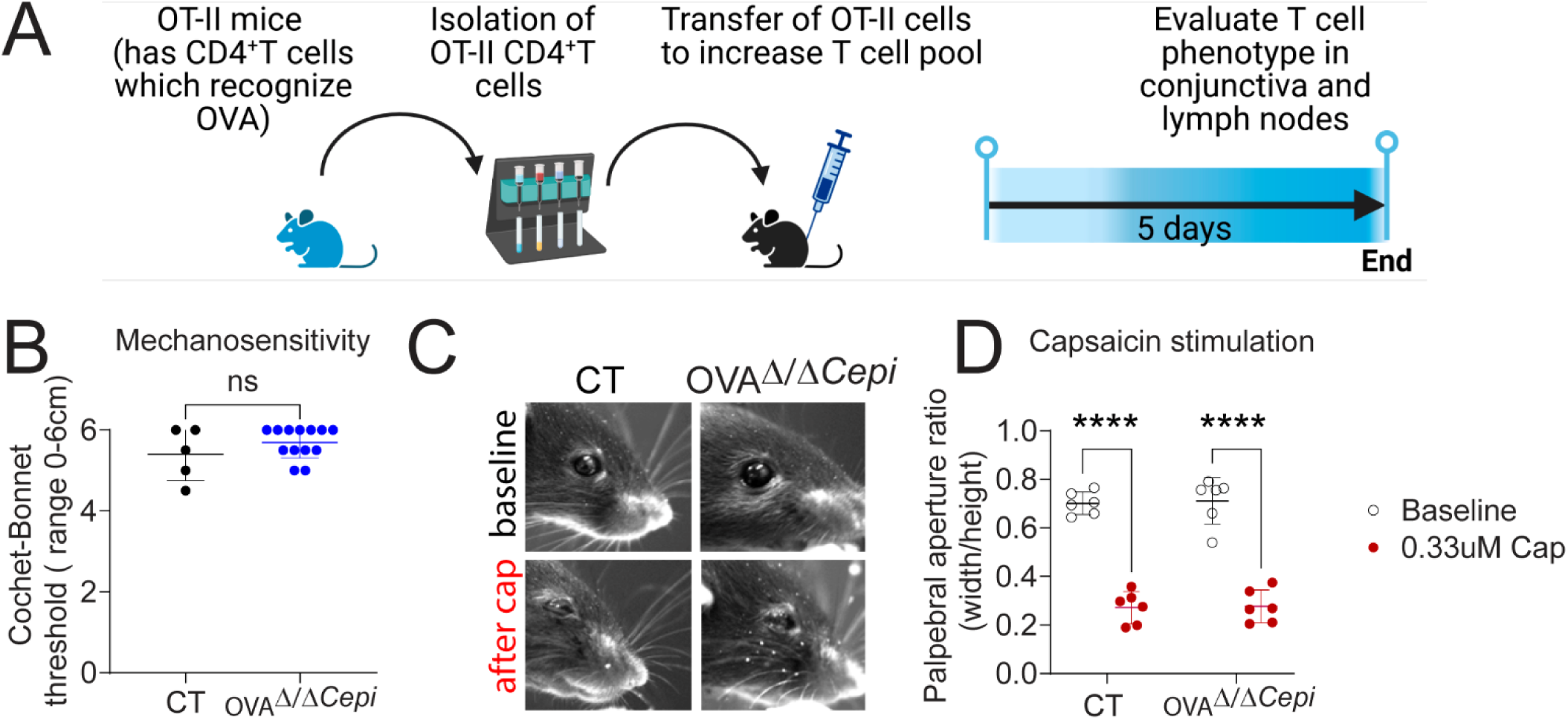
Neoantigen expression does not cause dry eye after adoptive transfer of OVA-specific T cells. **A.** Schematic of adoptive transfer of OT-II cells. OT-II cells were magnetically selected and adoptively transferred via intraperitoneal injection into control or OVA*^Δ/ΔCepi^* mice. Mice were euthanized 5 days post transfer. **B.** Mechanosensitivity measured with Cochet-Bonnet esthesiometer 2 weeks after dietary induction (range 0-6) and 5 days post adoptive transfer. High values are expected in naive mice. **C-D**. Functional assay using capsaicin stimulation as described in methods. Representative still images are shown in **C** and cumulative data in **D**. CT = control. ****P<0.0001, ns = non-significant

Next, we interrogated the fate of the corneal antigen-specific CD4^+^T cells by harvesting the ocular draining lymph nodes and conjunctiva 5 days after adoptive transfer. We applied multidimensional (19-marker) flow cytometry analysis of the whole CD4^+^T cell population in these samples with the intent to not only identify corneal antigen-specific cells, but also to assess if they assumed an activation, regulatory or anergic/exhaustion phenotype. The panel included antibodies for naïve (CD44^-^CD62L^+^), regulatory (CD4^+^CD25^+^Foxp3^+^), effector or cytokine-producing (IL-17A^+^IFN-γ^+^) (Chauhan et al., 2009; El Annan et al., 2009; Thomann et al., 2021), anergic (Foxp3^-^CD44^+^CD73^+^FR4^+^CTLA-4^+^Egr-2^+^CD25^+^), and activated (CD69^+^, CD25^+^, CD44^+^, and CD71^+^) (El Annan et al., 2009; Motamedi et al., 2016). OT-II cells are trackable based on their transgenic Vα2/Vβ5 T cell receptor. First, we measured their frequency in ocular draining nodes and observed a significant increase in CD45^+^CD3^+^CD4^+^Vα2^+^Vβ5^+^ cells in OVA*^Δ/ΔCepi^* mice (1.6±0.7 vs. 2.5±0.8%, p=0.035, Fig 5A). While there was not an overall increase in total CD4^+^Foxp3^+^CD25^+^ cells (Fig 5B), there was a significant increase in Vα2^+^Vβ5^+^ cells within the CD4^+^CD25^+^Foxp3^+^ population in the OVA*^Δ/ΔCepi^* group compared to controls (Fig. 5C-E). Within the activated non-Treg cells (CD4^+^CD25^+^Foxp3^-^), the frequency of Vα2^+^Vβ5^+^ did not change (Fig. 5E). There was also a detectable increase in the overall frequency of IFN-γ-producing CD4^+^T cells, but not within the Vα2^+^Vβ5^+^ cells (Fig. 5F-H), and no change in the frequency of IL-17-producing CD4^+^T cells (Fig. 5I-K). In the conjunctiva, we observed no change in the frequency of CD45^+^CD3^+^CD4^+^Vα2^+^Vβ5^+^ (Supplemental Fig. 2). Interestingly, we observed a decrease in total CD4^+^Foxp3^+^CD25^+^ cells (Supplemental Fig. 2), but no change in the OVA-specific Tregs. Similarly to the draining nodes, there was no change in frequency of activated non-Treg OVA-specific T cells nor of IFNγ/IL-17-producing CD4^+^T cells (Supplemental Fig. 2).

**Figure 5.**
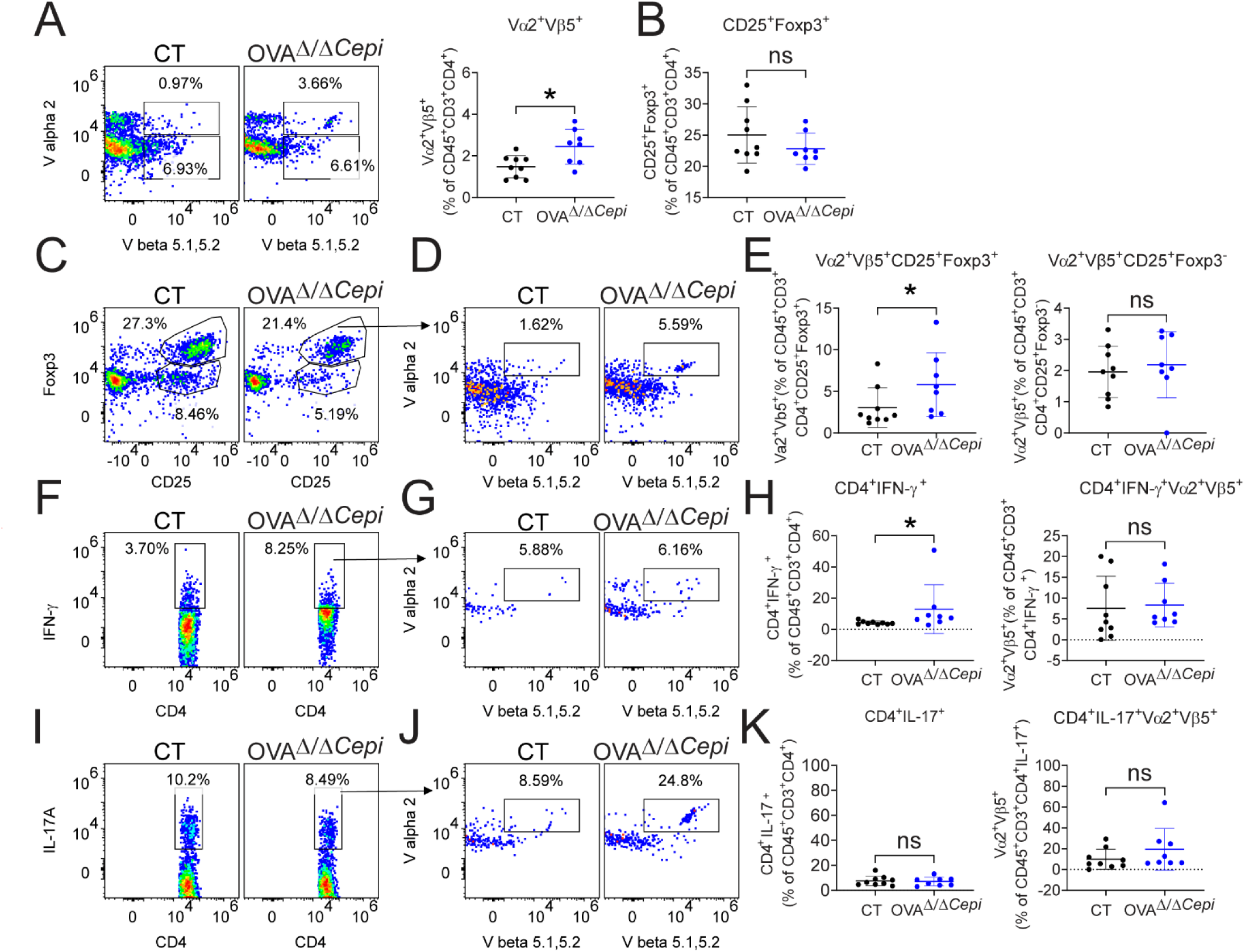
T cell fate after encountering a corneal antigen for the first time in ocular draining lymph nodes. OT-II cells were magnetically selected and adoptively transferred intraperitoneally into control or OVA*^Δ/ΔCepi^* mice. Mice were euthanized 5 days post transfer and ocular draining lymph nodes were excised and prepared for flow cytometry. CD45^+^CD3^+^CD4^+^T cells were gated and the frequency of Vα2^+^Vβ5^+^ and CD25^+^Foxp3^+^ cells were calculated within the total CD4^+^ population. Sequentially, the frequency of Vα2^+^Vβ5^+^ cells was calculated within the CD25^+^Foxp3^+^ and CD25^+^Foxp3^-^ populations. The frequency of Th1 and Th17 cells within the total CD4+T cells was calculated and the frequency of Vα2^+^Vβ5^+^ cells within these populations was also calculated. **A.** Representative dot plots and cumulative data of Vα2^+^Vβ5^+^ (gated on CD45^+^CD3^+^CD4^+^). **B.** Cumulative data of CD25^+^Foxp3^+^ (gated on CD45^+^CD3^+^CD4^+^). **C.** Representative dot plots of CD25^+^Foxp3^+^ and CD25^+^Foxp3^-^ cells. **D.** Representative dot plots identifying Vα2^+^Vβ5^+^ cells within the CD25^+^Foxp3^+^ population. **E.** Cumulative data of Vα2^+^Vβ5^+^ cells within CD25^+^Foxp3^+^ and CD25^+^Foxp3^-^ populations. **F-H**. Representative dot plots showing CD4^+^IFN-γ^+^ cells (**F**), the Vα2^+^Vβ5^+^ cells within this population (**G**), and cumulative graphs (**H**). **I-K**. Representative dot plots showing CD4^+^IL17^+^ cells (**I**), the Vα2^+^Vβ5^+^ cells within this population (**J**) and cumulative graphs (**K**). CT = control. * p<0.05; ns = non-significant. Mann-Whitney *U* test. Each dot represents a different animal.

Next, we performed unsupervised clustering of the combined CD4^+^ T cells from the conjunctiva (1,040 cells) and oDLNs (54,267 cells) according to their flow cytometry profiles. This analysis identified the 10 different populations shown in Fig. 6A. Despite the lower number of cells, the same populations but in different proportions could be identified in both tissues and across treatments (Figure 6B-C and Table 1). Figs 6D and 6E summarize the expression of each marker across the different CD4^+^T clusters. Populations 1-5 corresponded to endogenous, non-OVA-specific (Vα2^-^Vβ5^-^) CD4^+^T cells at different activation stages, as they were all positive for early growth response protein 2 (EGR2). EGR2 is a transcription factor induced by TCR signaling that is also associated with T cell anergy and exhaustion (Wagle et al., 2021). Of these, only cluster 1, which included CD4^+^T cells positive for the earliest markers of T cell activation (CD62L^-^, EGR2^+^, CD44^lo^), was expanded in OVA*^Δ/ΔCepi^* mice (Table 1). Population 6 comprised non-OVA-specific central memory Th17 cells (CD62LhiCD44hiIL-17A+), and population 7 comprised non-OVA-specific Tregs (Foxp3+CD25+ CD357+ CD152+); the proportions of both did not differ significantly between CT and OVA*^Δ/ΔCepi^* mice. Population 8 corresponded to the adoptively transferred OVA-specific (Vα2^+^Vβ5^+^) CD4^+^T cells, and as expected, these corneal antigen-specific cells were not the most abundant population. In addition to the Vα2^+^Vβ5^+^ TCR signature, this group was positive for Treg markers and IL-17A expression. There was a non-significant trend toward increased representation of this population in OVA*^Δ/ΔCepi^* mice, consistent with the predefined analysis (Fig. 5). Populations 9 and 10 comprised mostly endogenous CD4^+^T cells whose TCR included either the Vα2 or the Vβ5 chains of the transgenic OT-II cells. Because these cells lacked both chains, they were most likely not OVA-specific. These two populations resembled clusters 2-3 in the expression levels of all other markers.

**Figure 6.**
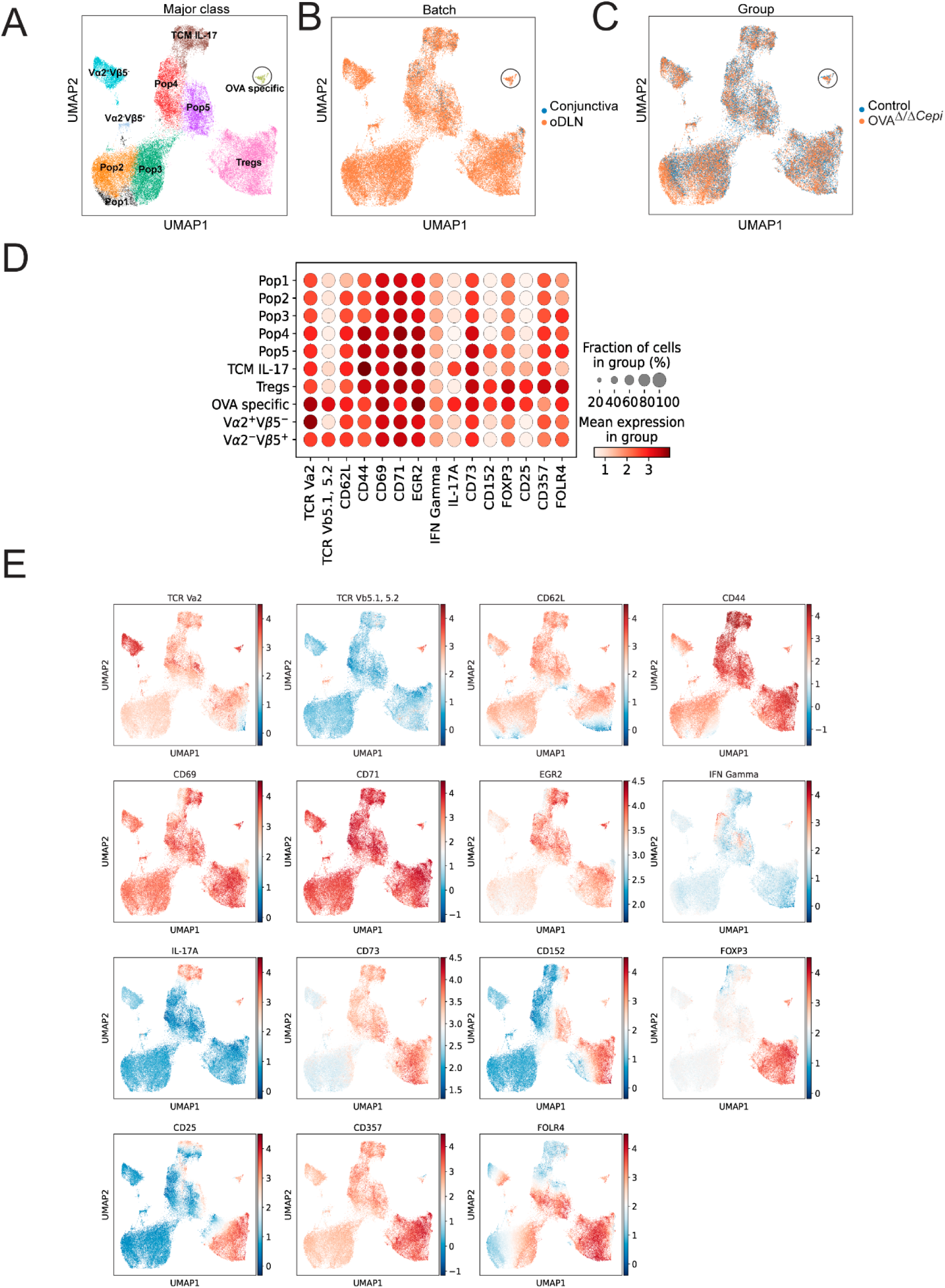
OVA specific T cells have a mixed phenotype. OT-II cells were magnetically selected and adoptively transferred intraperitoneally into control or OVA*^Δ/ΔCepi^* mice. Mice were euthanized 5 days post transfer and ocular draining lymph nodes were excised and prepared for flow cytometry. Unsupervised analysis of conjunctiva and ocular draining lymph nodes (oDLN) was performed as described in the Methods. **A-C**. UMAP visualization (**A**) of 10 identified CD4^+^T cell populations. OVA-specific cells are indicated within the black circle. UMAP visualizations colored by tissue origin (**B**) and experimental group (**C**). The uniform distribution of cells across clusters demonstrates successful batch effect removal and sample integration. **D.** Dot plot illustrating the mean expression level (color) and fraction of cells (size) for markers across the identified populations. **E.** UMAP feature plot indicating expression of markers within the different populations.

**Table 1.** Proportion of CD4^+^T cell clusters between control and OVA^Δ/ΔCepi^ groups. CD4^+^T cell clusters were identified using unsupervised clustering of high-dimensional flow cytometry data. The markers assessed were: Foxp3, CD25, CD73, CD357, CD69, CD71, FR4, EGR2, IL-17, CD62L, CD152, CD44, IFN-γ, TCR Vα2, and TCR Vβ5.

| CD4<sup>+</sup> T cell clusters | CLN |  |  | CJ |  |  |
| --- | --- | --- | --- | --- | --- | --- |
|  | Control<br>(%,<br>mean±SD) | OVA<sup>Δ/ΔCepi</sup><br>(%,<br>mean±SD) | Wilcoxon p | Control<br>(%,<br>mean±SD) | OVA<sup>Δ/ΔCepi</sup><br>(%,<br>mean±SD) | Wilcoxon p |
| Pop 1 | 0.26±0.23 | 7.63±9.57 | **0.00076** | 0.87±1.44 | 0.46±1.31 | 0.56 |
| Pop 2 | 16.59±2.66 | 14.23±4.99 | 0.39 | 3.12±2.33 | 5.89±5.49 | 0.23 |
| Pop 3 | 17.86±2.39 | 19.70±3.57 | 0.15 | 2.87±2.73 | 4.56±3.42 | 0.413 |
| Pop 4 | 12.07±1.77 | 9.71±1.14 | **0.0071** | 22.31±9.26 | 29.55±9.51 | 0.067 |
| Pop 5 | 9.67±2.09 | 7.69±1.67 | 0.054 | 23.67±5.89 | 21.98±7.49 | 0.66 |
| TCM IL-17 | 8.46±4.08 | 7.06±3.80 | 0.34 | 22.44±6.28 | 15.28±9.64 | 0.067 |
| Tregs | 28.02±4.46 | 24.73±2.27 | 0.10 | 22.28±8.54 | 19.28±7.50 | 0.56 |
| OVA specific | 0.63±0.65 | 0.95±0.91 | 0.50 | 1.14±1.48 | 0.90±1.42 | 0.74 |
| Vα2<sup>+</sup>Vβ5<sup>-</sup> | 5.68±0.84 | 7.31±0.65 | **0.0028** | 1.31±1.45 | 1.44±1.84 | 0.96 |
| Vα2<sup>-</sup>Vβ5<sup>+</sup> | 0.75±0.21 | 1.00±0.32 | 0.12 | 0.00±0.00 | 0.67±1.53 | 0.39 |

Altogether, our results indicate that most corneal antigen-specific CD4^+^T cells become Tregs upon encountering their cognate antigen in the eye-draining lymph nodes, and that this process is accompanied by increased naïve CD4^+^T cell recruitment from the blood and possibly also transient activation of non-corneal antigen-specific CD4^+^T cells (Weaver et al., 2009).

### After systemic immunization with adjuvants, cornea neoantigen expression leads to dry eye

In other organs such as the liver, pancreas, and bladder, *de novo* expression of a harmless foreign antigen tricks the immune system into treating the introduced antigen as self-derived and like other autoantigens in the corresponding tissue (Cebula et al., 2013; Legoux et al., 2015; Liu et al., 2007; Riehn et al., 2017). Our findings so far indicate that, under homeostatic conditions, corneal neoantigen expression induces tolerance via Treg induction in the ocular-draining lymph nodes. However, established peripheral tolerance may be lost if antigen-specific CD4^+^ T cells are primed elsewhere (Horai et al., 2015; Salaman and Gould, 2020), thus leading to organ-specific disease. To study this possibility, we first switched OVA*^Δ/ΔCepi^* and control mice to a doxycycline diet for 2 weeks, allowing tolerance to corneal epithelial OVA neoexpression to develop (as in Fig. 3). Then, the newly established immune balance was challenged by subcutaneous injection. immunization with OVA emulsified in complete Freund’s adjuvant as a source of potent innate immune activators, and its outcome was read two weeks later by a delayed-type hypersensitivity (DTH) assay, a clinically validated surrogate of the antigen-specific effector immune response (Guzman et al., 2014; Ko et al., 2018). Indeed, previously-tolerant-but-later-immunized OVA*^Δ/ΔCepi^* mice exhibited a full DTH response comparable to non-tolerized-and-later-immunized control mice **(Fig. 7B)**. Since this indicated that immunization in both groups led to the development of an OVA-specific effector immune response (overcoming previously established tolerance to the same corneal epithelium-expressed antigen in OVA*^Δ/ΔCepi^* mice), we then examined the eyes. We hypothesized that the OVA-specific CD4^+^T cells would encounter the corneal antigen and trigger autoimmunity. We observed a non-significant trend toward decreased corneal mechanosensitivity in OVA*^Δ/ΔCepi^* mice (**Fig. 7C**), suggesting OVA-driven effector CD4^+^T cell activity, and increased corneal epithelial staining in this group, which confirmed barrier disruption via a corneal epithelium-targeting immune response (**Fig. 7D**). Altogether, these findings indicate that peripheral immune tolerance to a corneal epithelial neoantigen is not absolute and may be subverted by strong innate immune activation, leading to autoimmunity.

**Figure 7.**
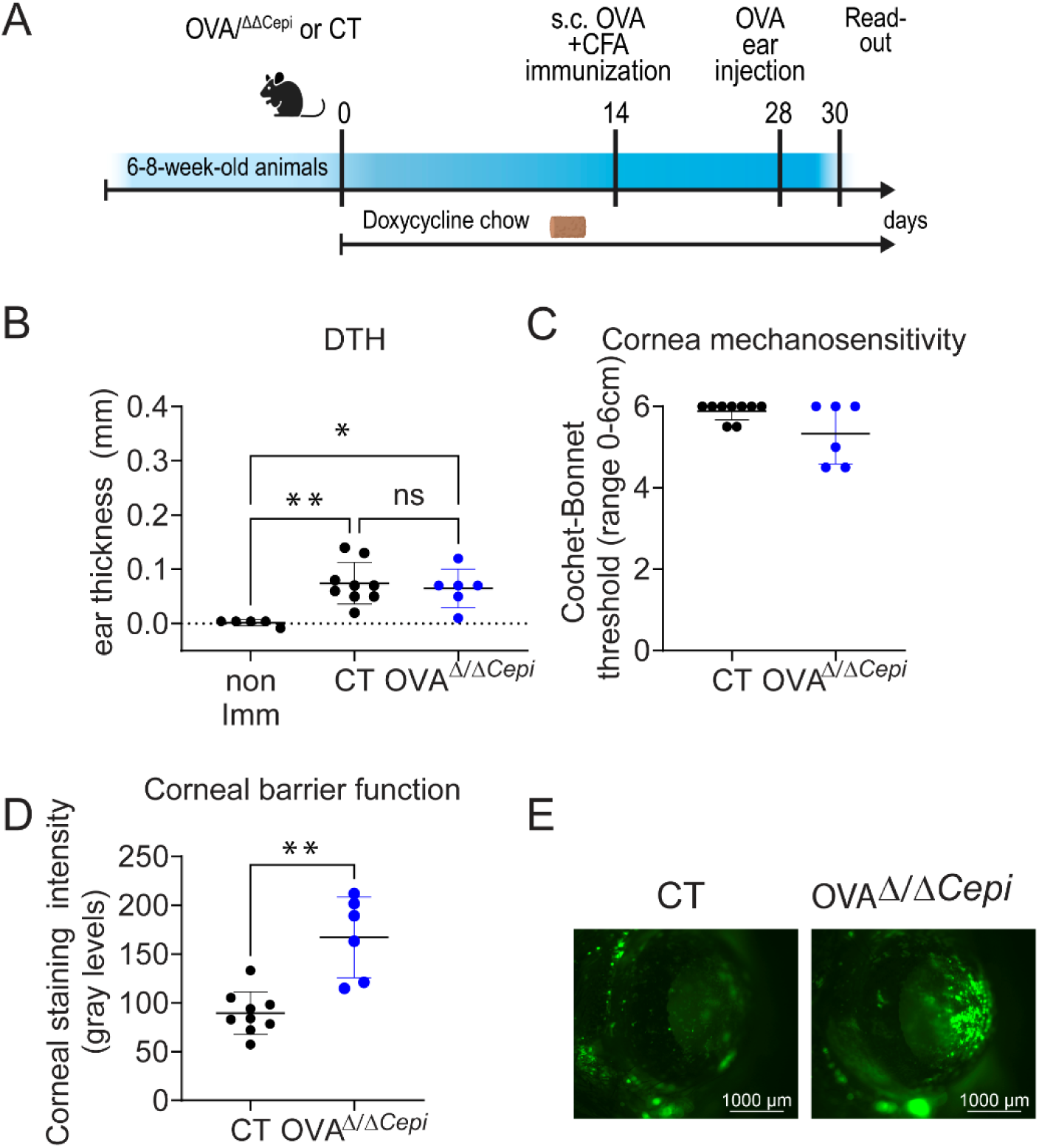
Immunization with OVA leads to dry eye phenotype. **A.** Experimental schematic. Mice received doxycycline chow 2 weeks before (day 0) the immunization with OVA and complete Freund’s adjuvant (CFA). On day 28 (four weeks after switch to doxycycline chow), mice received an intradermal OVA injection in the ear. Ear swelling was measured 48 hours post-injection. Ear swelling in immunized mice was compared to non-immunized mice (non-imm). **B.** Ear swelling 2 weeks post-immunization. Kruskal-Wallis with Dunn’s posthoc test, n = 4-9 mice/group. Imm = immunization with Complete Freund’s adjuvant and OVA. **C.** Mechanosensitivity measured with Cochet-Bonnet esthesiometer. High values are expected in control mice. **D-E.** Corneal barrier function was measured as the uptake of a fluorescent dye. Cumulative data (**D**) and representative images (**E**) are shown. Higher numbers indicate worse corneal barrier function. CT = control. Mann-Whitney *U* test. n = 6-9. Each dot represents a mouse. *P<0.05; **P<0.01; ns = non-significant.

## Discussion

It is widely accepted that CD4^+^T cells play a key pathogenic role in dry eye and other ocular surface disorders that involve the cornea(Chen et al., 2014; Niederkorn et al., 2006; Vereertbrugghen et al., 2024; Vereertbrugghen et al., 2023). While significant progress has been made in understanding the basis of corneal immune privilege and successful allogeneic corneal transplantation(Lee et al., 2025; Niederkorn, 2015), the regulatory mechanisms that prevent immune responses against corneal epithelial antigens and ocular surface inflammation remain incompletely understood(Galletti and de Paiva, 2021; Galletti et al., 2017). In this study, we developed an inducible mouse model that enables the controlled expression of a neoantigen restricted to the corneal epithelium, allowing us to use antigen-specific CD4^+^T cells to track the immune response to this antigen. We found that the *de novo* expression of a corneal epithelial antigen is actively tolerated under homeostatic conditions by mechanisms that differ from those underlying corneal immune privilege. Instead of relying on ACAID or antigen sequestration, corneal epithelial immune homeostasis is maintained primarily by inducing antigen-specific Tregs in the ocular-draining lymph nodes. However, this tolerance is not absolute, as it can be overcome by innate immune activation and replaced by the expansion of pathogenic effector T cells that target the cornea and induce epithelial damage. Thus, our results shed light on the immune mechanisms underpinning ocular surface homeostasis and disease at the antigen-specific cell level for the first time.

Many self-antigens are expressed in the thymus, leading to central tolerance by deletion of potentially autoreactive T cells or their differentiation to natural Tregs before entering the mature T cell pool (Meng et al., 2023). However, this process only applies to the fraction (∼85% of the coding genome) (Danan-Gotthold et al., 2016) of self-antigens expressed in the thymus and does not encompass all the autoantigens found systemically or newly generated antigens. For the remaining non-thymically-expressed autoantigens, peripheral tolerance serves as an additional, tissue specific protective layer (ElTanbouly and Noelle, 2021; Legoux et al., 2015; Malhotra et al., 2016). Antigens expressed exclusively in the intestinal or bronchial epithelium (but not during thymic education) lead to the expansion of specific Tregs in the periphery, i.e., the draining lymph nodes. By contrast, pancreatic β cell-restricted antigens do not expand Tregs, and their cognate CD4^+^T cells remain ignorant of their expression. In the case of interphotoreceptor retinoid-binding protein (a well-known retinal autoantigen in uveitis), T cells specific for one epitope are deleted in the thymus while those reactive against another one within the same protein are not (Taniguchi et al., 2012). Until now, the lack of models allowing antigen-specific tracking of T cells responding to corneal epithelial antigens prevented understanding ocular surface-specific immune tolerance (Galletti and de Paiva, 2021; Galletti et al., 2017). By combining the ROSA26OVA system with corneal epithelial-specific inducible Cre activity, we generated a ternary mouse model in which ovalbumin is expressed selectively in the corneal epithelium. We confirmed that upon induction, the antigen was processed and presented physiologically: by corneal epithelial cells on MHC I molecules, and by ocular surface-derived antigen-presenting cells (APCs) on both MHC I and MHC II molecules, as the latter could stimulate OVA-specific CD4^+^T cell proliferation *in vitro* (Fig. 2). The absence of lymphatic vessels in the non-inflamed cornea has been highlighted as a critical factor underlying tolerance via antigen sequestration/immunological ignorance, particularly in corneal transplantation (Dietrich et al., 2010; Niederkorn and Larkin, 2010; Weaver et al., 2009). Conversely, allogeneic corneal grafting into a vascularized bed considerably increases the risk of rejection, partly because it allows CCR7-mediated trafficking of corneal dendritic cells to the eye-draining lymph nodes (Jin et al., 2007). Our findings indicate that corneal epithelial antigens are immunologically accessible under non-inflammatory conditions and can be captured and presented by eye-draining APCs. By contrast, ACAID relies on bloodborne F4/80^+^ macrophages that transport antigens from the anterior chamber to the spleen (Lin et al., 2005). It remains to be established whether corneal-resident APCs directly migrate to lymph nodes or if conjunctival APCs take up shed corneal epithelial cells, as is the case for topically delivered or exogenous antigens (Galletti et al., 2013; Ko et al., 2018). This observation effectively rules out strict immunological ignorance as the principal tolerance mechanism for corneal epithelial antigens and instead supports the existence of active peripheral tolerance mechanisms.

In our model, neoantigen expression in the cornea did not cause spontaneous ocular pathology (Fig. 3), consistent with tolerance. Mice expressing corneal OVA did not exhibit changes in corneal epithelial barrier function or mechanosensitivity thresholds upon antigen induction nor after prolonged antigen expression. While the system was designed for OVA expression restricted to corneal epithelial cells, the abundant intraepithelial nerves are highly sensitive to immune activity in the ocular surface and would thus have evidenced abnormal T cell activation. Moreover, antigen expression alone did not activate APCs in ocular draining lymph nodes, suggesting that corneal antigen presentation occurs in the absence of co-stimulatory signals under homeostatic conditions. These findings align with the concept that peripheral tolerance mechanisms operate in tissues as long as antigens are continuously presented without accompanying danger signals (Belkaid and Oldenhove, 2008; Mueller, 2010; Salaman and Gould, 2020). Consistently, the frequency of OVA-specific CD4^+^T cells increased in the eye-draining lymph nodes of mice expressing corneal OVA, indicating that these cells recognized their cognate antigen *in vivo* and confirming that immunological ignorance was not at play. A significant fraction (but not all) of these antigen-specific cells acquired a regulatory phenotype (Foxp3, CD25, and CTLA-4 expression), while there was no increase in activated non-regulatory antigen-specific T cells or evidence of Th1/Th17 polarization (Fig. 6). These findings suggest that the dominant outcome of antigen encounter in this context is the differentiation of antigen-specific Tregs rather than effector cells. Remarkably, corneal OVA expression also led to an increase in non-adoptively transferred CD4^+^T cells in the ocular draining lymph nodes, which were presumably non-OVA-specific and expressed markers consistent with TCR engagement. We speculate that this corresponds to abortive T cell activation in response to epitope spreading and competition for the immunodominant OVA-derived peptide recognized by the adoptively transferred OT-II cells (Weaver et al., 2009), as there was no observable ocular phenotype.

Our results explain how the corneal epithelium maintains immune tolerance despite constant exposure to environmental stressors and potential epithelial damage. The ocular surface is continuously subjected to desiccation, microbes, and mechanical stress, all of which can trigger innate immune pathways (Bron et al., 2017; Chi et al., 2017; Redfern et al., 2013; Reins et al., 2018; Yu et al., 2021a; Yu et al., 2023). Yet under physiological conditions, corneal inflammation is kept at bay (Barabino et al., 2012; Galletti et al., 2017). Our results indicate that peripheral induction of antigen-specific Tregs in ocular draining lymph nodes is key to maintaining immune homeostasis against corneal epithelial antigens. These findings differ from the reported immune privilege mechanisms that mediate tolerance in corneal transplantation, which involve splenic presentation of endothelial antigens and sequestration of corneal epithelial and stromal antigens due to lack of lymphatic vessels (Chen et al., 2022; Hori et al., 2019; Taylor, 2016). Also, ocular exposure to an exogenous antigen can induce conjunctival mucosal tolerance via uptake by conjunctival APCs and the generation of Tregs. By contrast, the tolerance mechanism described here involves *de novo* expression of a tissue-restricted antigen within the corneal epithelium itself. Our results therefore suggest the existence of an additional layer of ocular immune regulation that specifically controls responses to endogenous corneal epithelial antigens, which is likely critical in dry eye pathogenesis.

Remarkably, tolerance to corneal epithelial antigens in this model was not absolute. When mice that had expressed and tolerated corneal OVA for weeks were immunized systemically with the same antigen along with an adjuvant, they developed a robust delayed-type hypersensitivity response and corneal epithelial barrier disruption (Fig. 6). These findings indicate that antigen-specific effector T cells generated under inflammatory conditions can override peripheral tolerance and target the cornea once they encounter their cognate antigen *in situ*. In other words, the corneal epithelium is not intrinsically resistant to immune-mediated injury, as its antigens are visible and accessible to the immune system. Instead, immune tolerance normally prevents the generation of pathogenic autoreactive T cells. This observation has potential implications for dry eye disease pathogenesis. Autoreactive CD4^+^T cells drive ocular surface inflammation in dry eye (Chen et al., 2014; Niederkorn et al., 2006; Vereertbrugghen et al., 2024; Zhang et al., 2011), though the antigens recognized by these cells remain unknown. Our findings suggest that, under physiological conditions, corneal epithelial antigens induce tolerance by generating antigen-specific Tregs. However, environmental stressors or inflammatory signals that activate APCs could shift this balance toward effector T cell priming, as has been shown for exogenous antigens (Guzman et al., 2016a; Guzman et al., 2016b; Guzmán et al., 2020; Guzman et al., 2018). Once generated, these pathogenic T cells could target the cornea and perpetuate the vicious inflammatory cycle underlying dry eye disease.

Our study has limitations. First, OVA represents a model antigen that does not necessarily mimic the biochemical properties of endogenous corneal proteins. Second, the adoptive transfer of OT-II T cells increases the precursor frequency of antigen-specific T cells beyond physiological levels, which could potentially influence tolerance mechanisms; however, experiments with a physiological T cell repertoire had the same physiological outcomes, making this possibility unlikely. Third, while our data indicate that antigen-specific Tregs are induced in ocular-draining lymph nodes, the specific APC subsets responsible for antigen presentation and Treg induction remain to be defined. More research is warranted on which dendritic cell populations mediate this process and how environmental stressors alter their function during ocular surface inflammation.

In summary, we demonstrated that a corneal epithelial neoantigen is actively tolerated rather than being ignored by the immune system. This form of peripheral tolerance is mainly mediated by Tregs generated in the ocular-draining lymph nodes and can be overcome by strong inflammatory priming. These findings provide a mechanistic framework for understanding how immune homeostasis is maintained at the ocular surface and, conversely, how its breakdown may contribute to autoimmune inflammation in dry eye disease.

### Disclosures

The authors declare no competing interests,

## Material and Methods

### Animals

The Institutional Animal Care and Use Committee at Baylor College of Medicine approved all animal experiments. In addition, all studies adhered to the Association for Research in Vision and Ophthalmology for the Use of Animals in Ophthalmic and Vision Research and the NIH Guide for the Care and Use of Laboratory Animals (National Research Council Committee for the Update of the Guide for the and Use of Laboratory, 2011).

### Creation of inducible, cornea-restricted neoantigen mouse model

Using the Cre-Flox system, we generated a ternary keratin 12-cornea-specific, doxycycline-inducible tetO-Cre, OVA-transgene line. We mated ROSA26OVA^f/f^ mice (Cebula et al., 2013; Legoux et al., 2015; Riehn et al., 2017) (obtained through an MTA from Dr. Wirth, Medical University Hannover, Germany) with Keratin 12-reverse tetracycline-trans-activator knock-in/tetO-cre (*Krt12^rtTA/rtTA^*/tetO-cre (Chikama et al., 2005), creating a mouse in which activation of Cre irreversibly leads to OVA expression in the corneal epithelium (hereafter referred to as OVA*^Δ/ΔCepi^*). The Kr12 mouse line was obtained from Dr. Winston Kao (University of Cincinnati, Cincinnati, OH) and it was bred with tetO-cre mice from Jackson Laboratories (STOCK Tg(tetO-cre)1Jaw/J, stock number 006224). Krt12^rtTA^ is a cornea-specific promoter(Chikama et al., 2005; Kao et al., 1996; Liu et al., 1993; Yoshida et al., 2006). The OVA-containing gene sequence is reversed (thus -OFF-under naïve conditions, **Fig. 1A**) and flanked by oppositely oriented loxP sites. Each transgene allele was identified by PCR genotyping (Transnetyx, Cordova, TN).

Cre recombinase expression, under the control of a tetracycline-responsive element, was induced in mice by replacing their regular chow with doxycycline chow (200mg/kg, Bio-Serv, Flemington, NJ) *ad libitum*. OVA*^Δ/ΔCepi^* mice were induced at post-natal day 30 or older for two weeks to induce the irreversible expression of OVA in the corneal epithelium. By feeding mice doxycycline chow (a synthetic tetracycline), Cre becomes activated, translocates to the nucleus, and inverts the OVA cassette, permanently inducing OVA expression in the corneal epithelium. Mice used for experiments received doxycycline chow for at least two weeks. A separate group of mice received doxycycline for 4 months (from 2 to 6 months of age). Double-transgenic mice (Kr12^rtTA/rtTA^:OVA^f/f^), littermates also received a doxycycline diet and were used as controls.

### OT-II mice and Adoptive Transfer

OVA-specific T cell receptor transgenic mice were purchased from Jackson laboratories (B6.Cg-Tg(TcraTcrb)425Cbn/J, stock 4194, Jackson Laboratories, Bar Harbor, ME) and were used as donors in adoptive transfer experiments and in vitro assays at 8-12 weeks of age. CD4^+^T cells were isolated from the spleen and lymph nodes of OT-II mice using magnetic beads (Miltenyi Biotec, Bergisch Gladbach, Germany) and adoptively transferred (2×10^6^/mouse) into the experimental groups. The switch to doxycycline chow coincided with the adoptive transfer, which turned ON the expression of OVA in the corneal epithelium of OVA*^Δ/ΔCepi^* mice. Mice were euthanized 5 days after adoptive transfer + antigen induction, and the type of immune response in T cells in the cornea, conjunctiva, and ocular draining nodes were investigated using flow cytometry, which are described below.

### RNA isolation and qRT-PCR4

Transgene expression in the corneal epithelium, conjunctiva, and ear skin was evaluated in 8-week-old mice with and without doxycycline induction (n = 5-7/group). Corneal epithelium was collected by scraping. The conjunctiva was surgically excised from the bulbar and palpebral area. Samples were immediately placed into an Eppendorf tube with guanidine-isothiocyanate–-containing lysis buffer followed by selective binding of RNA to the silica–gel–based membrane (RNeasy Micro kit Plus, add catalog #; Qiagen, Gaithersburg, MD). The RNA concentration was measured by its absorption at 260 nm, and the samples were stored at -80°C until used for polymerase chain reaction (PCR). First-strand cDNA was synthesized from 0.5 µg of total RNA with random hexamers by M-MuLV reverse transcription (Ready-To-Go You-Prime First Strand Beads; GE Health Care, Inc., Arlington Heights, IL, catalog #27926401). Real-time PCR was performed with specific primers for GAPDH (CTCCCACTCTTCCACCTTCG and CCACCACCCTGTTGCTGTAG) and ovalbumin-fusion protein (CAGGCACTCCTTTCAAGACC and GCGGTTGAGGACAAACTCTT, specific for the ‘ON state’) with PowerTrack SYBR Green master mix (Applied Biosystems/ThermoFisher, Waltham, MA, catalog #A46110), in a commercial thermocycling system (QuantStudio 3D Digital PCR system (Life Technologies, Carlsbad, CA), according to the manufacturer’s recommendations. Assays were performed in duplicate in each experiment. The *Gpadh* gene was used as an endogenous reference for each reaction. The real-time PCR results were analyzed using the comparative CT method. The relative mRNA level in the ear skin was used as the calibrator.

### Whole Mount Cornea Immunostaining

Whole mount corneas (n = 4-5/group) were prepared as previously published (Vereertbrugghen et al., 2023) and stored at -80 °C until ready for use. Corneas were stained with an anti-ovalbumin antibody (MilliporeSigma, Merck KGaA, Darmstadt, Germany, catalog #AB1225), washed, and incubated with Alexa Fluor® 488 AffiniPure® Goat Anti-Rabbit IgG (H+L) antibody (Jackson ImmunoResearch, West Grove, PA, catalog #111-545-003) and Hoechst 33342 nuclei staining. (ThermoFisher Scientific, catalog # H3570). Whole corneas were flattened on microscope slides and covered with antifade medium (Vectashield Plus Antifade Mounting Medium, Vector, Newark, CA, catalog #H-1900-2), and coverslips were applied. Wholemount digital Images (512 x 512 pixels) were captured using a laser scanning Nikon confocal microscope (Nikon A1 RMP, Nikon, Melville, NY) with a wavelength of 400–750 nm. Whole-thickness images were captured using a 0.5 µm Z-step (whole-mount corneas only). The images were processed using NIS-Elements AR 6.10.01 (Nikon)

### Corneal Mechanical Sensitivity

Corneal sensitivity was measured with the Luneau Cochet-Bonnet Instrument as previously described using the step technique (Yu, 2022). Briefly, this instrument increases pressure as the filament shortens (6 cm to 0.5 cm). A clear stimulus-evoked blink and retraction of the eye into the ocular orbit indicated a positive response. The central cornea was tested six times at each filament length. The response was considered negative when the monofilament touch failed to elicit a blink. A positive response was considered when the animal blinked for 50% or more of the time tested. Since this test was performed in multiple different setups, the final sample size is described in the figure legend.

### Video Imaging of Palpebral Aperture Dimensions Analysis

Palpebral aperture size was assessed using a custom video-based imaging approach adapted from previously published methods (Yaman et al., 2025). Mice were gently restrained in a 50 mL conical tube that stabilized the body while allowing natural head positioning and unobstructed visualization of the ocular surface. Video recordings were acquired using an Alvium 1800 U-511m monochrome camera (Allied Vision, Stadtroda, Germany) equipped with a 25 mm fixed focal length C-Series lens (f/1.4), mounted at a constant distance from the animal to maintain consistent image scaling across experiments. Videos were captured at 60 frames per second for 2 min using StreamPix 9 software (NorPix, Montreal, Canada). To enhance contrast and reduce background variability, a matte black backdrop was placed behind the animals, and all recordings were performed at the same imaging station under standardized lighting conditions.

Each mouse was first recorded under unstimulated baseline conditions and then recorded again following topical application of capsaicin (0.33 µM). Experimental cohorts consisted of OVA*^Δ/ΔCepi^* mice and littermate controls (n = 6/group). Both groups received doxycycline chow for at least 2 weeks before the experiment. Video files were reviewed using frame-by-frame inspection, and one representative frame was selected approximately every 100 frames for analysis. If the designated frame was compromised by motion or focus artifacts, a neighboring frame within a ±10-frame range was used instead. Palpebral aperture dimensions were quantified manually using ImageJ (NIH, Bethesda, MD, USA) by measuring eyelid height and eyelid width in pixels. To account for minor variations in camera positioning or animal alignment, aperture size was expressed as a height-to-width ratio (HWR), yielding a normalized, dimensionless metric for comparisons across genotypes and experimental conditions. All recordings were conducted at similar times of day to minimize circadian variability.

### Antigen Presenting Cell Functional Assay

As previously reported (Bian, 2018), single-cell suspensions of cervical lymph nodes from OVA and control mice (n = 4-5/group) were prepared (de Souza et al., 2021), filtered with 100µm CellTrics filters (Sysmex, Kobe, Japan; catalog # 04-004-2328), incubated with mitomycin C (50 μg/ml for 1 h at 37 C followed by two 10-minute-long wash/incubation steps in complete media) and used as antigen presenting cells (APC) in vitro. OT-II CD4^+^T cells were selected using a magnetic isolation kit (Miltenyi) as described above, incubated in Cell Violet Tracer (Invitrogen, ThermoFisher, catalog #C34571) and used as responding T cells. APC (100,000cells/100μl) and responding T cell suspensions (100,000 cells/100μl) were co-plated with and without added OVA (100 μg/ml; Sigma-Aldrich, St. Louis, MO) in RPMI 1640 media (Gibco, ThermoFisher, catalog #61870-036) supplemented with 10% fetal bovine serum, 50ug/ml of gentamicin and 1.25ug/ml of amphotericin B. Plates were photographed daily and cells were collected at day 4 for proliferation assays using violet tracer dilution flow cytometry experiments. A BD LSRII Benchtop cytometer was used for data acquisition, and data was analyzed using BD Diva Software (BD Pharmingen, San Diego, CA) and FlowJo software (version 10.1; Tree Star, Inc., Ashland, OR). Biological replicates were averaged.

### Flow Cytometry Analysis of Cornea, Ocular Draining Lymph Nodes and Conjunctiva

For investigation of antigen expression in the cornea, OVA*^Δ/ΔCepi^* and control mice were used (n = 9/group) after at least two weeks on doxycycline chow. The corneas were excised (two per mouse) and transferred to a solution of 20 mM EDTA for 20 minutes. Corneas were washed with PBS twice, before being placed in a solution of 0.1% collagenase IV (Gibco ThermoFisher; catalog #17104-019) in Hank’s Balanced Salt Solution (Gibco; catalog #14025-092), chopped, and incubated at 37°C for thirty minutes. DNAse I (MilliporeSigma; catalog #260913-10mu) was added to each suspension in the last fifteen minutes. Suspensions were neutralized with RPMI (Gibco; catalog #61870-036) and filtered with 100µm CellTrics filters (Sysmex; catalog #04-004-2328) before proceeding with staining. Single-cell suspensions were incubated with CD16/32 (clone 2.4G2; BD Biosciences, Franklin Lakes, New Jersey; catalog #553142) to block Fc receptors for ten minutes on ice before being stained with an infrared fluorescent reactive live/dead dye diluted 1:3000 (Invitrogen ThermoFisher; catalog #L34993) for fifteen minutes. Cells were then fixed with a 2% formaldehyde solution (FisherScientific ThermoFisher; catalog #BP531-25) for twenty minutes, before being stained overnight with the following antibodies: CD45_BV510 (clone 30-F11; BioLegend, San Diego, California; catalog #103138), CD326-APC (EPCAM) (clone G8.8; BioLegend; catalog #118214), SIINFEKL-PE (clone 25-D1.16; BioLegend; catalog #141603), CD11b-PECy7 (clone M1/70; BioLegend; catalog #101216), and MHC II-PerCP eFluor 710 (clone AF6-120.1; Invitrogen; catalog #46-5320-82). Suspensions were washed four times the next day before being acquired on a BD Canto II Benchtop cytometer with BD Diva software version 6.7 (BD Biosciences). The final data was analyzed using FlowJo software version 10 (Tree Star).

To investigate T cell fate after adoptive transfer of OT-II cells, OVA*^Δ/ΔCepi^* and control mice were used (n = 7-8/group) after at least two weeks on doxycycline chow. As previously reported, single-cell suspensions of cervical lymph nodes were prepared (de Souza et al., 2021) and filtered with 100µm CellTrics filters (Sysmex; catalog #04-004-2328) before proceeding with staining. Conjunctivae were excised, digested in collagenase as previously published (Bian, 2018) and made into single cell suspensions. 1×10^6^ cells from ocular draining lymph nodes and 100µl of conjunctival single cell suspensions were plated in a 96-well U bottom plate and then stained. Single-cell suspensions were incubated with CD16/32 (clone 2.4G2; catalog #553142) to block Fc receptors for ten minutes on ice before being stained with an infrared fluorescent reactive live/dead dye diluted 1:3000 (Invitrogen ThermoFisher; catalog #L34993) for fifteen minutes.

According to the manufacturer’s protocol, cells were fixed and permeabilized with BD Pharmingen Transcription Buffer Set (BD Biosciences; catalog #562574). Cells were then stained overnight with the following: FOLR4-PE Dazzle 594 (clone 12A5; BioLegend; catalog #125015), CD73-BV605 (clone TY/23; BioLegend; catalog #117205), IFN-γ-BV480 (clone XMG1.2; ThermoFisher; catalog #414-7311-82), CD62L (L-Selectin)-BV510 (clone MEL-14; BioLegend; catalog #104441), CD71-PerCP-Cy5.5 (clone RI7217; BioLegend; catalog #113815), T cell receptor Vβ5.1, 5.2-BV750 (clone MR9-4; BD BioSciences; catalog #746897), EGR2-PE (clone erongr2; Invitrogen; catalog #12-6691-82), CD4-FITC (clone RM4-5; Cytek Biosciences, Fremont, CA; catalog #35-0042-U500), CD69-BV711 (clone H1.2F3; BioLegend; catalog #104537), CD25-PE Fire 700 (clone PC61; BioLegend; catalog #102081), IL-17A-BV650 (clone 17B7; ThermoFisher; catalog #416-7177-82), CD357-BV421 (clone DTA-1; BioLegend; catalog #126331), CD152 (CTLA-4)-PE

Fire 640 (clone UC10-4B9; BioLegend; catalog #106333), Foxp3-PE Cy7 (clone 3G3; Cytek Biosciences; catalog #60-5773-U100), CD44-PerCP Fire 806 (clone IM7; BioLegend, catalog #103081), T cell receptor Vα2-BUV 661 (clone B20.1; BD Biosciences; catalog #750116), CD45-Alexa Fluor 700 (clone 30-F11; BioLegend; catalog #103128), CD3-APC Fire 810 (clone 17A2; BioLegend; catalog #100267), and CD8-Spark Blue 574 (clone 53-6.7; BioLegend; catalog #100794). Suspensions were washed four times the next day before being acquired on a Cytek Aurora spectral flow cytometer with SpectraFlo software (Cytek). The final data was analyzed using FlowJo software version 10 (Tree Star).

### High Dimensional Flow Cytometry Data Processing

Compensated flow cytometry data were exported from FlowJo (BD Biosciences) and imported into the cyCONDOR package (v0.3.1)(Kroger et al., 2024) for initial processing. Data were imported and auto-logically transformed using the prep_fcs() function to normalize signal intensities. Dimensionality reduction was performed by first computing Principal Component Analysis (PCA) via the runPCA() function, followed by Uniform Manifold Approximation and Projection (UMAP) using the runUMAP() function to facilitate low-dimensional visualization. The normalized expression matrices, associated metadata, principal components, and UMAP coordinates were exported and integrated into Scanpy (v1.11.2) (Wolf et al., 2018) for downstream analysis. Unsupervised clustering was performed using the Leiden algorithm (scanpy.tl.leiden). Cell clusters and gene expression profiles were visualized using UMAP embeddings (scanpy.pl.umap). Gene expression patterns across clusters were further characterized using dot plots (scanpy.pl.dotplot). To evaluate differences in cell type proportions between the control and OVA groups, the Wilcoxon rank-sum test was applied. Statistical significance was defined as a p-value < 0.05.

### Measurement of Corneal Barrier Function

Corneal barrier function was assessed by quantifying corneal epithelial permeability to 70-kDa AlexaFluor®OregonGreen Dextran (OGD; Invitrogen ThermoFisher, catalog #D7172), according to a previously published protocol (Yu et al., 2021b) in OVA*^Δ/ΔCepi^* and control mice after either two weeks or four months of doxycycline chow. Briefly, 1 µL of a 50 mg/mL OGD solution was applied to the ocular surface one minute before euthanasia, which was performed using excess isoflurane followed by cervical dislocation. Corneas were rinsed with 2 mL of PBS and photographed with a stereoscopic zoom microscope (model SMZ 1500; Nikon) under fluorescence excitation at 470 nm. OGD staining intensity was quantified in digital images by measuring the mean fluorescence intensity within a 2-mm diameter circle placed on the central cornea using NIS Elements software (version AR, 5.20.02). This assessment was conducted independently by two observers. The mean intensity of the right and left eyes was averaged, and then the resulting mean from biological replicates was calculated and analyzed. Since this test was performed in multiple different set-ups, the final sample size is described in the figure legend.

### Delayed-Type Hypersensitivity (DTH) Assay

OVA*^Δ/ΔCepi^* (n = 6) and control mice (n = 9) were switched to doxycycline chow for 14 days before the immunization (day 0). Mice received doxycycline chow for the duration of the experiment. On day 14, mice were immunized with an emulsion of OVA and complete Freund’s adjuvant (ThermoFisher) prepared at a 1:1 ratio and administered subcutaneously in the neck under general anesthesia using isofluorane gas dispensed through a nose cone with the SomnoSuite Vaporizer (Kent Scientific, Torrington, CT). On day 28, mice were challenged with the OVA antigen by intradermal ear injection (10 μg of OVA in the right ear). Ear swelling was measured after 48 hours using a gauge micrometer (Mitutoyo, Kanagawa, Japan). The range of swelling in the injected ear was within range of our previously published studies (Barbosa et al., 2017; Ko et al., 2018). A group of naive unimmunized mice (n = 5) served as controls.

## Statistical Analysis

Based on pilot studies, the sample size was calculated with StatMate2 Software (GraphPad Software, San Diego, CA). Statistical analyses were performed with GraphPad Prism software (GraphPad Inc, CA, version 9.2). Data were first evaluated for normality with the Kolmogorov-Smirnov normality test. Appropriate parametric (t-test) or non-parametric (Mann-Whitney) statistical tests were used to compare the two age groups. Whenever adequate, one-way or two-way ANOVA or Kruskal-Wallis followed by post hoc tests were used. The final sample per experiment is shown in the legends.

## Figure Legends

**Supplemental Figure 1 (relates to Figure 2):**
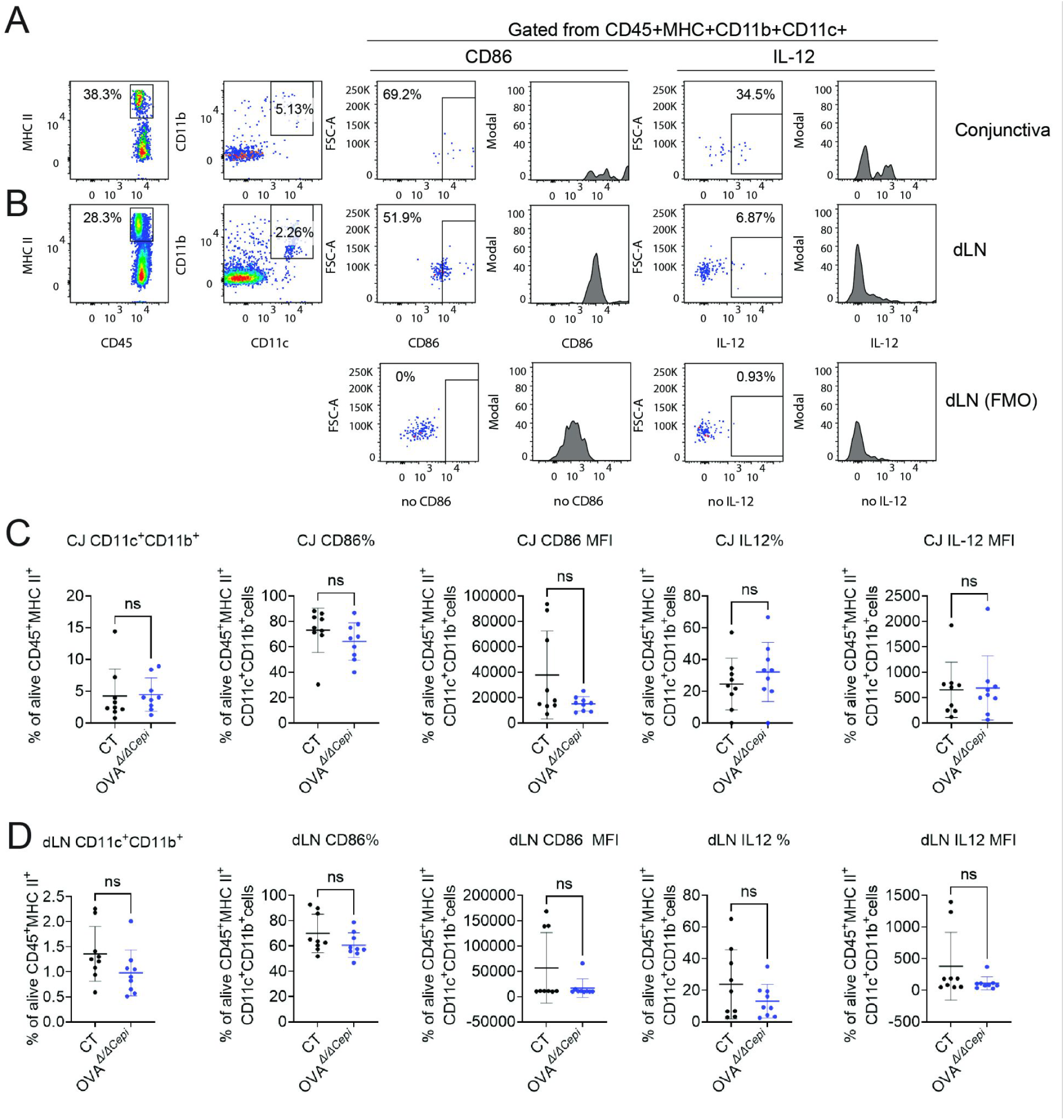
APCs are not activated after neoexpression of OVA in the cornea. After dietary induction, conjunctiva and lymph nodes were collected and processed for flow cytometry to evaluate CD86 and IL-12 expression. **A-B**. Gating strategy in conjunctiva (**A)** and ocular draining lymph nodes (dLN) (**B**). **C-D**. Cumulative data showing frequency of DCs in conjunctiva (**C**) or ocular draining lymph nodes (**D**). Mean±SD, each dot represents an animal. Mann-Whitney U test, ns = non-significant. CT = control.

**Supplemental Figure 2 (relates to Figure 5).:**
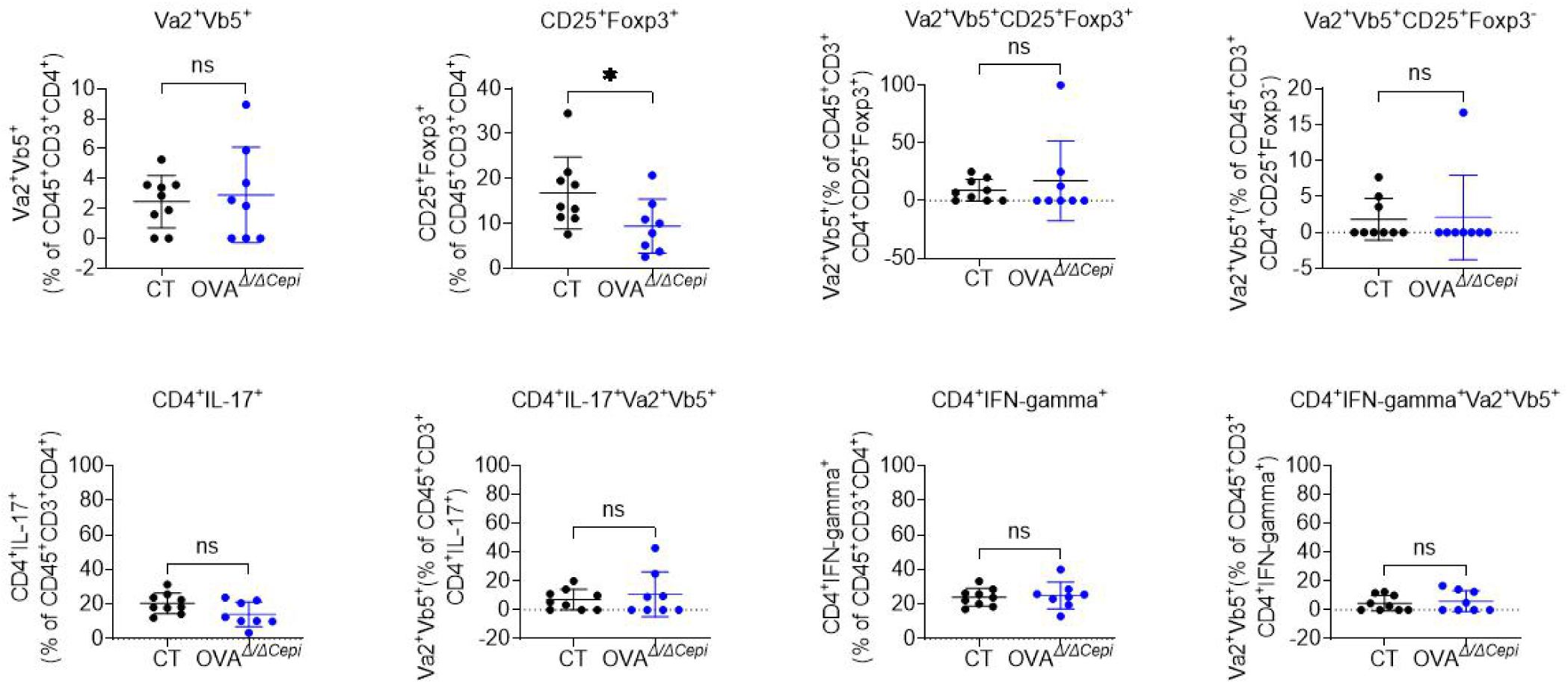
T cell fate after encountering a corneal antigen for the first time in conjunctiva. OT-II cells were magnetically selected and adoptively transferred via intraperitoneal injection into control or OVA*^Δ/ΔCepi^* mice. Mice were euthanized 5 days post transfer and conjunctivae were excised and prepared for flow cytometry. CD45^+^CD3^+^CD4^+^T cells were gated, and the frequency of Vα2+Vβ5+ and CD25+Foxp3+ cells was calculated within the total CD4^+^T population. Sequentially, the frequency of Vα2^+^Vβ5^+^ cells was calculated within the CD25^+^Foxp3^+^ and CD25^+^Foxp3^-^ populations. **A-D.** Cumulative data of Vα2^+^Vβ5^+^ (**A**, gated on CD45^+^CD3^+^CD4^+^), CD25^+^Foxp3^+^ (**B**, gated on CD45^+^CD3^+^CD4^+^) and Vα2^+^Vβ5^+^CD4^+^CD25^+^Foxp3^+^ **(C)** cells and Vα2^+^Vβ5^+^CD4^+^CD25^+^Foxp3^-^ cells (**D**). Mann-Whitney *U* test. Each dot represents a different animal. *P<0.05; ns = non-significant. CT = control.

## Notes

**Support:** This work was supported by the Sjögren Foundation, NIH EY026893 (CSDP); NIH/NEI EY002520 (Core Grant for Vision Research Department of Ophthalmology); NEI Training Grant in Vision Sciences T32 EY007001 (KKS); BCM Genomic & RNA Profiling Core GARP Core [P30 Digestive Disease Center Support Grant (NIDDK-DK56338) and P30 Cancer Center Support Grant (NCI-CA125123), NIH S10 grant (1S10OD02346901)]. Further research support was provided by Research to Prevent Blindness (unrestricted grant to the Department of Ophthalmology), The Hamill Foundation, and The Sid Richardson Foundation. Jeremias Galletti received a Fulbright Visiting Scholar Award to participate in this study, and he is funded by Wellcome Trust 221859/Z/20/Z and Agencia Nacional de Promoción Científica y Tecnológica (Argentina, PICT 2020-00138, PICT 2021-00109).

### Competing Interest Statement

The authors have declared no competing interest.

